# Evolution of Auxin Response Factors in plants characterized by phylogenomic synteny network analyses

**DOI:** 10.1101/603175

**Authors:** Bei Gao, Liuqiang Wang, Melvin Oliver, Moxian Chen, Jianhua Zhang

## Abstract

Auxin response factors (ARFs) have long been a research focus and represent a class of key regulators of plant growth and development. Previous studies focusing genes from limited number of species were unable to uncover the evolutionary trajectory of this family. Here, more than 3,500 ARFs collected from plant genomes and transcriptomes covering major streptophyte lineages were used to reconstruct the broad-scale family phylogeny, where the early origin and diversification of *ARF* in charophytes was delineated. Based on the family phylogeny, we proposed a unified six-group classification system for angiosperm ARFs. Phylogenomic synteny network analyses revealed the deeply conserved genomic syntenies within each of the six ARF groups and the interlocking syntenic relationships connecting distinct groups. Recurrent duplication events, such as those that occurred in seed plant, angiosperms, core eudicots and grasses contributed the expansion of ARF genes which facilitated functional diversification. Ancestral transposition activities in important plant families, including crucifers, legumes and grasses, were unveiled by synteny network analyses. Ancestral gene duplications along with transpositions have profound evolutionary significance which may have accelerated the functional diversification process of paralogues. Our study provides insights into the evolution of ARFs which will enhance our current understandings for this important transcription factor family.

## INTRODUCTION

The plant hormone auxin, the chemically simple molecule indole-3-acetic acid, controls many physiological and developmental processes in land plants including but not limited to organogenesis, tissue differentiation, apical dominance, gravitropism as reviewed previously (Kieffer *et al.*, 2010). Completion of the genome of the moss *Physcomitrella patens* makes revealed that many core functional proteins required for auxin biosynthesis, perception, and signaling are present in this early-diverging land plant (embryophyte) lineage (Rensing *et al.*, 2008, Lang *et al.*, 2018), and suggested the auxin molecular regulatory network evolved at the latest in the common ancestor of mosses and angiosperms. A recent review suggested that with the exception of the PGP/ABCB auxin transporters, homologues of all the other core components for hormonal control of physiology and development by auxin could be identified in *P. patens* (Thelander *et al.*, 2018). Changes in auxin perception and signaling that occurred through evolution could have generated the diversification of plant forms that occurred during the past ~474-515 million-year history of the land plants (Morris *et al.*, 2018), which eventually led to the complex vegetative innovations that shape the modern terrestrial and freshwater ecosystems.

Auxin response factors (ARF), as core components in auxin signaling, have long been a focus of plant signaling research (Chapman and Estelle, 2009). The twenty-three *ARFs* identified in the *Arabidopsis thaliana* genome were phylogenetically clustered into three subfamilies (Clades A, B and C) which were subsequently divided into seven groups (ARF9, ARF1, ARF2, ARF3/4, ARF6/8, ARF5/7 and ARF10/16/17); a classification that was well supported by *ARF* genes from other angiosperms and representative non-flowering lineages (Finet *et al.*, 2013). Generally, ARF proteins can be functionally divided into transcriptional activators (ARF5-8 and 19 in *A. thaliana*) and repressors (remaining ARFs in *A. thaliana*) with well-characterized functional domain architectures (Guilfoyle and Hagen, 2007, Finet *et al.*, 2013). ARFs bind to the auxin response elements (AuxRE: TGTCTC) in the promoter region of downstream auxin-inducible genes (Ulmasov *et al.*, 1997) and function in combination with Aux/IAA repressors, which dimerize with ARF activators in an auxin-regulated manner (Ulmasov *et al.*, 1999, Guilfoyle and Hagen, 2007). Unlike ARF activators, few reports have demonstrated that ARF repressors are able to interact with other ARF proteins or Aux/IAA proteins (Vernoux *et al.*, 2011). A recent top-notch work revealed a newly identified mechanism where the IAA32 and IAA34 transcriptional repressors are stabilized by the transmembrane kinase 1 (TMK1) at the concave side of the apical hook to regulate ARF gene expression and ultimately inhibit growth (Cao *et al.*, 2019).

In most of the well-established transcription factor annotation procedures, such as those implemented by the PlnTFDB(Perez-Rodriguez *et al.*, 2010), PlantTFDB(Jin *et al.*, 2017), iTAK(Zheng *et al.*, 2016) and TAPScan(Wilhelmsson *et al.*, 2017), ARFs were identified using two signature domains: the B3 (PF02362) domain and the Auxin-resp (PF06507) domain, although some ARF proteins (e.g. ARF23 in *A. thaliana)* may be truncated and lack the Auxin-resp domain(Guilfoyle and Hagen, 2007). Finet *et al.* established a robust and comprehensive phylogenetic framework for the ARF gene families(Finet *et al.*, 2013), however *ARF* genes from non-flowering plants were under-represented. Comprehensive annotation of transcription factors covering distinctive plant clades demonstrated that a number of plant specific transcription factor families (including ARF) evolved in streptophytic algae (charophytes)(Wilhelmsson *et al.*, 2017), suggesting an earlier origin of ARF than that proposed by Finet and collegues(Finet *et al.*, 2013).

Compared to conventional gene family studies that focus on one or a limited number of species of interest(Kalluri *et al.*, 2007, Wang *et al.*, 2007, Wang *et al.*, 2012b), phylogenetic studies on a broader scale that include multiple plant lineages were able to generate a more robust insights into the evolutionary process that gave rise to the modern assemblage of a target gene family (Finet *et al.*, 2013, Li *et al.*, 2015). The inclusion of genomic synteny data provides important information that impacts the determination of the evolutionary past of a gene family, especially when the gene family of interest evolved in parallel with ancestral genome duplication events (Gao *et al.*, 2018). The conventional genomic block alignment that connects orthologues, retained on genomic syntenic blocks, worked well for a limited number of species(Cheng *et al.*, 2013, Gao *et al.*, 2018), but a network approach was more effective when multiple genomes were included in the synteny analyses(Zhao *et al.*, 2017, Zhao and Schranz, 2017). A comprehensive genomic synteny network can be constructed using nodes to represent the target genes and associated adjacent genomic blocks and the network edges (connecting lines) to represent syntenic relationships(Zhao and Schranz, 2017, Gao *et al.*, 2018). The recently established phylogenomic synteny network methodology was able to integrate and summarize genomic synteny relationships to uncover and place genomic events (e.g. ancient tandem duplications, lineage-specific transposition activities) into the evolutionary past of a target gene family(Zhao *et al.*, 2017, Zhao and Schranz, 2017).

In this study, we collected more than 3,500 ARF members to generate a comprehensive gene-family phylogeny with the aim of filling evolutionary gaps in the non-flowering plants and splitting the long branches present in the current phylogeny(Finet *et al.*, 2013). We propose an updated model for evolution of the ARF family that covered the major streptophytic clades that was based on the six-group classification system we proposed for the ARF genes in angiosperms. A phylogenomic synteny network analyses of angiosperm genomes revealed the deep positional conservation of *ARF* gene-family members within each of the six groups. Detailed individual synteny network analyses together with phylogenetic reconstructions for the six ARF groups revealed their distinctive evolutionary histories. Ancestral duplication events in angiosperms, and subsequent WGDs in eudicots and monocots have contributed to the expansion of ARF members. Ancestral lineage-specific transpositions in important angiosperm families such as crucifers, legumes and grasses were also unveiled. Together, the results presented here add to our current understanding of the evolutionary process that established *ARF* genes in plants. We also expect this broad-scale evolutionary framework could help direct future functional studies that further explore the interplay between auxin signaling and the evolution of land plants.

## RESULTS AND DISCUSSION

### Auxin Response Factor Evolved in Streptophytic Alage

To generate a broad-scale phylogenetic profile for ARF genes in plants, we collected a total of 3,502 *ARF* homologues in the streptophytes (including charophytes and embryophytes). *ARFs* were present in all major clades of streptophytes including charophytes, hornworts, liverworts, mosses, lycophytes, ferns and seed plants (Supplementary Fig. S1). In chlorophytes, the Auxin-resp domain was not detected although some chlorophyte genes did contain the B3 domain. Genes containing both the B3 (PF02362) and the Auxin_resp (PF06507) domains were identified in streptophytic algae (charophytes). This was consistent with the observation that a number of plant-specific transcription factors evolved in streptophytic algae (Wilhelmsson *et al.*, 2017). Charophytes represented a paraphyletic clade encompassing successive sister lineages to the land plants (Leliaert *et al.*, 2012, Wilhelmsson *et al.*, 2017). We identified *ARF* homologues in species that are found in three charophyte orders: Zygnematales, Coleochaetales and Chlorokybales, but *ARF* homologues were not identified in the transcriptomes of Charales and Klebsormidiales. However, the presence of ARF in charophytes was affirmed by the *Chara braunii* (Charales) genome (Nishiyama *et al.*, 2018). The identification of the single *ARF* gene in *Chlorokybus atmophyticus* (Chlorokybaceae) suggests that the origin of ARF genes traces back to the root position of streptophytes (Supplementary Fig. S1). This observation suggests an earlier origin of *ARF* gene than those reported previously (Finet *et al.*, 2013, Wilhelmsson *et al.*, 2017).

### Broad-scale Phylogenetic Profile of ARFs in plants

Overall, the numbers of *ARF* genes in individual angiosperm genomes are greater than those in the individual genomes of non-flowering plants and the ‘recent’ polyploids, such as *Glycine max,* possess conspicuously more *ARF* genes than other plants (Supplementary Fig. S2). The inclusion of homologues identified from the 1KP transcriptome database provided a comprehensive atlas for the ARF family phylogeny. Overall, the broad-scale phylogeny of *ARFs* generated in this analysis was closely in parallel with the phylogenetic relationships among plant lineages (Fig. 1) derived from large-scale phylotranscriptomic study(Wickett *et al.*, 2014). The phylogenetic tree generated from the ARF gene collection was rooted by the *ARF* gene from *Chlorokybus atmophyticus,* an early diverging charophyte, and exhibited a consistent tree topology with that reported previously(Finet *et al.*, 2013). The incorporation of transcriptomic data from non-flowering plants enabled long evolutionary branches to be split. The phylogenetic analyses also provided robust evidence that angiosperm *ARFs* could be separated clearly into three major subfamilies (Clade A, B and C; consistent with previously reported groupings(Finet *et al.*, 2013)). The three major subfamilies encompassed the six groups (designated as Group I through VI) in this study (Fig. 1). In comparison to the classification proposed by Finet *et al*.(Finet *et al.*, 2013), the Group-I *ARFs* contained the ARF1 and ARF9 subfamilies (which were likely to have derived from an ancient angiosperm-wide duplication) and Group II through VI correspond to the ARF 2, ARF 3/4, ARF 6/8, ARF 5/7 and ARF 10/16/17 subfamilies (Supplementary Table S1), respectively.

**Figure 1.**
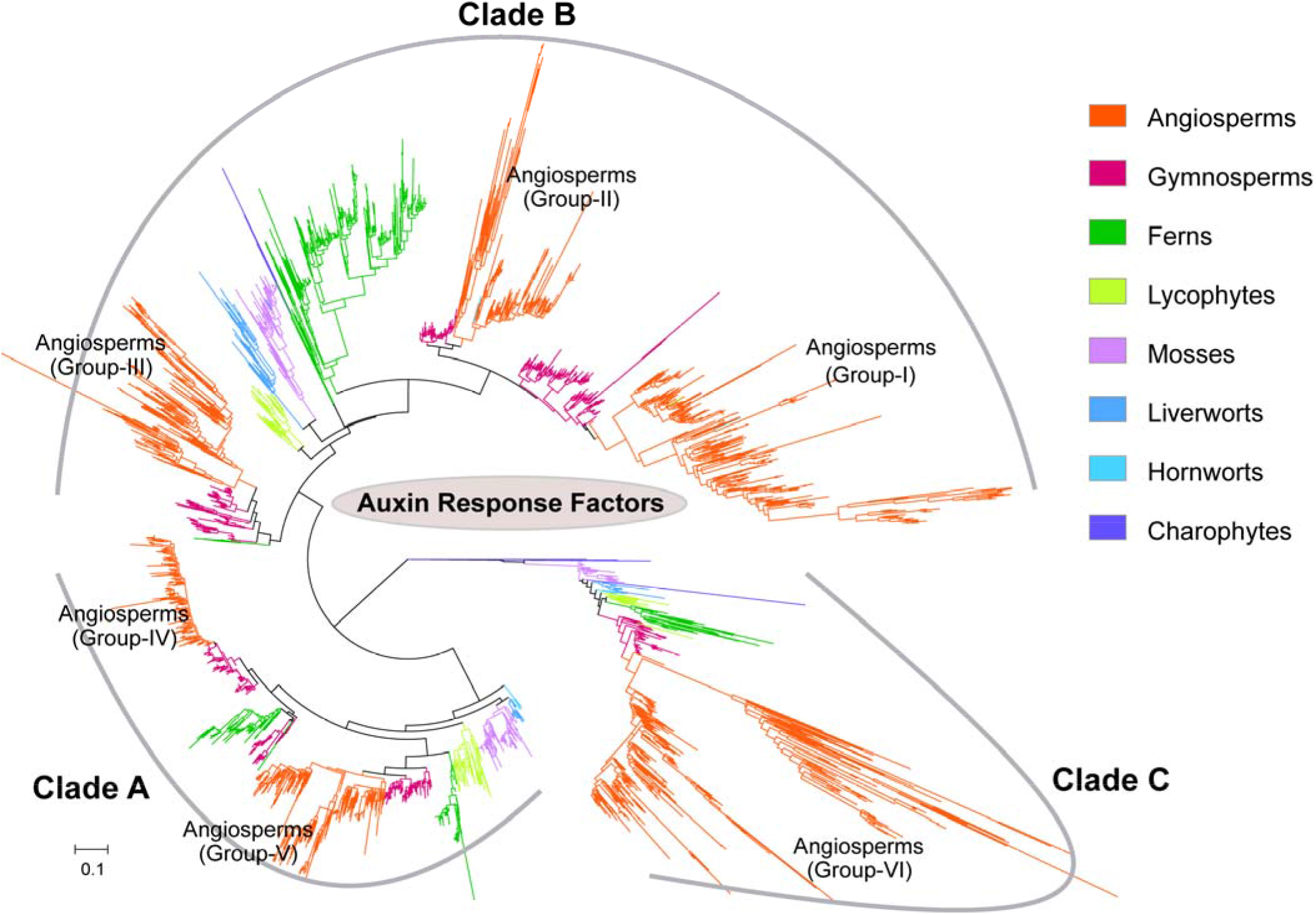
The broad-scale family phylogeny of *ARF* genes in plants. The broad-scale family phylogeny of ARF genes in different plant lineages estimated using IQ-TREE maximum-likelihood algorithm. Branches representing *ARF* genes from different plant clades were colored and six conspicuous groups for angiosperm *ARF* genes were obtained and labeled on the tree (Groups I through VI). Gene tree structure and the three subfamilies (denoted as clades A, B and C) were consistent with that reported in (Finet *et al.* 2013) with more *ARF* genes identified from plant genomic or transcriptomic datasets.

Groups I, II and III were clustered in the subfamily Clade-B, Groups IV and V in Clade A and Group Vi in Clade C. Clade C was revealed to be basal and a sister clade to subfamilies A plus B. *ARF*s from the charophytes (Zygnematales and Coleochaetales) were separated into clade-C and clade-B failing to partition into a basal mono- or para-phyletic clade, which suggested an ancient diversification of *ARF* genes within the charophytes. The family phylogeny also revealed that each of the six angiosperm ARF family groups was located with gymnosperm *ARF* genes as the closest sister lineage. The tree branches of gymnosperm *ARF* genes are conspicuously shorter than those for angiosperms (Fig. 1), which suggested lower amino acid substitutional rates and higher levels of protein sequence conservation in gymnosperm *ARF* genes likely a result of longer generation times that are common in the gymnosperms(Smith and Donoghue, 2008).

Clade-A contains *ARF* genes that cover all major embryophyte (land plant) clades and contains ARF genes of group-III together with orthologues from gymnosperms and ferns. The *ARF* genes from seed plants and ferns were separated into two major clades which are sister to each other which resulted in a tree topology that was consistent with two child clades derived from an ancient duplication. While lycophyte *ARF* genes were placed outside of and sister to the large duplication clade shared by ferns and seed plants. *ARF* genes from hornworts were identified as basal-most in Clade-A, followed by genes from mosses and liverworts.

Clade-B was the most diversified lineage containing the angiosperm group I and II genes and along the gymnosperm orthologous genes delineated a conspicuous seed-plant duplication (the *ζ* event)(Jiao *et al.*, 2011). However, ARF genes from hornworts, liverworts and ferns were mixed into this large duplication clade (Fig. 1). We hypothesize that they might be derived from convergent evolution, though the possibilities of horizonal gene transfer or sequence contaminations cannot be eliminated. Genes from ferns, mosses, liverworts and lycophytes were placed as successive sister lineages to this duplication clade.

Clade-C was situated as the basal clade with a relatively simple phylogenetic profile and contains genes from every major plant lineage (from charophytes to angiosperms, Fig. 1). This configuration updated the evolutionary model in which clade-C *ARFs* were absent in gymnosperms(Finet *et al.*, 2013).

The broad-scale phylogenetic analyses in this study established a robust and unified six-group classification system for angiosperm *ARF* genes, which is consistent with previous phylogenetic and domain architecture studies(Guilfoyle and Hagen, 2007, Finet *et al.*, 2013). The relative phylogenetic positions of other land plant lineages were also clarified (Fig. 1), providing a consistent phylogenetic framework for subsequent synteny network analyses.

### Evolutionary trajectory of ARFs augmented with current genomic and transcriptomic data

In concordance with the phylogenetic analyses described by Finet *et* al.(Finet *et al.*, 2013), we augmented the evolutionary trajectory of ARF family in plants with gene sequences from the currently available genomic and transcriptomic data. The resulting phylogenetic trajectory path (Fig 2.) suggested that the three ARF subfamilies (clades A, B and C) were likely diversified through an ancient duplication in the charophytes, which is consistent to the evolutionary trajectory proposed previously (Flores-Sandoval *et al.*, 2018, Mutte *et al.*, 2018). Tree uncertainties and unresolved land plant phylogenies were also reflected in the ARF gene-family phylogeny, leaving some of the evolutionary processes elusive.

**Figure 2.**
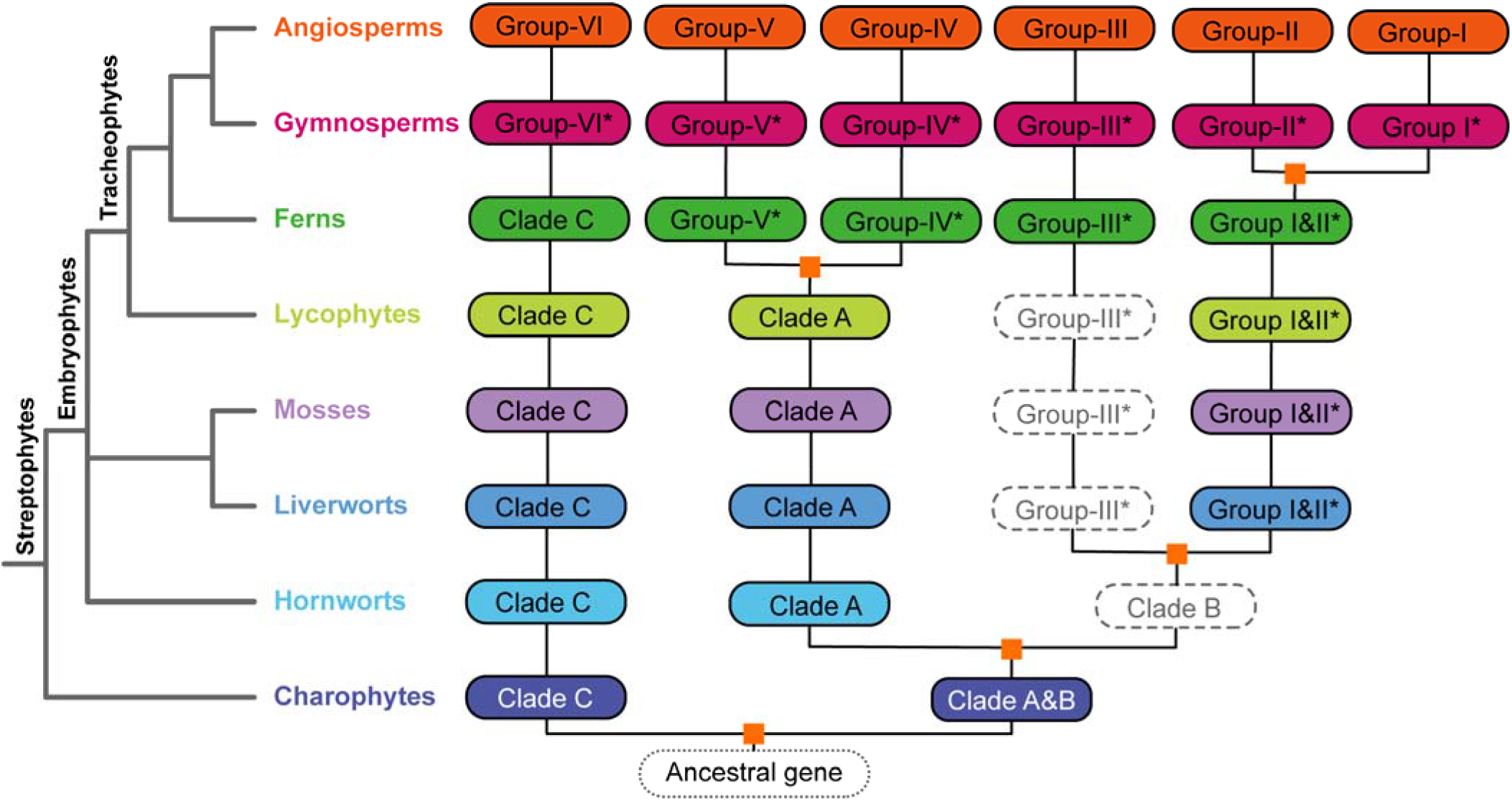
Ancient evolutionary trajectory of *ARF* genes in streptophytes. Annotation of auxin response factor genes suggested an early origin and diversification in streptophytic algae. The eight major plant lineages were represented with different colors, solid rounded rectangles indicate presence of ARF genes in corresponding plant lineages, dashed ones suggested absence of data and potential gene losses. Gene groups indicated with an asterisk represented close sister lineages to corresponding angiosperm *ARFs*. Inferred ancient gene duplications were depicted as golden squares. Group-I and Group-II ARFs was suggested to be derived from the seed plant duplication event.

All of the ARF transcriptional activators (ARF 5-8 and 19 in *A. thaliana)* were clustered in the clade-A subfamily. Within clade-A, the ARF genes were well-conserved in all land plant lineages and appear to have experienced a conspicuous ancient duplication event that occurred in the ancestor of ferns and seed plants. This ancient duplication generated groups IV (ARF6 and 8 in *A. thaliana)* and V (ARF5, 7 and 19 in *A. thaliana)* in the angiosperms and the corresponding sister groups in gymnosperms and ferns (Fig. 2). The *ARF* genes in bryophytes (including hornworts, liverworts and mosses) and lycophytes (clubmosses) were outside of this duplication. The *ARF* genes in clade-A also exhibited a gene tree topology consistent with the ‘hornwort-sister’ land plant phylogeny in contrast to the ‘bryophytes-monophyletic’ phylogeny(Wickett *et al.*, 2014, Morris *et al.*, 2018). The evolutionary well-conserved aspect of the ARF activator genes indicates an early genetic foundation for auxin signaling networks in the embryophytes(Thelander *et al.*, 2018).

In clade-B, unlike clade-A, *ARF* genes from hornworts were not found densely populated at the basal position of the subtree and some hornwort *ARFs* were found clustered with angiosperm *ARF* genes, making the evolution of clade-B ARFs in hornworts elusive. The trajectory analysis suggests two ancient duplications in clade-B, an embryophyte duplication shared by mosses, liverworts and tracheophytes that occurred before the diversification of groups I/II and group III, and a seed plant duplication that generated the groups: I and II. However, the close sister groups for group III were only found in the gymnosperms and ferns which suggested there were gene losses in mosses, liverworts and lycophytes (Fig. 2).

The subfamily of clade-C is well-conserved, covering all streptophytic lineages, and generated the simplest phylogenetic profile (Fig. 2), containing the group-VI angiosperm *ARF* genes (the ARF 10, 16, and 17 in *A. thaliana).* Hornwort *ARF* genes were placed as direct sisters to the vascular plants (tracheophytes) and the *ARF genes of* mosses and liverworts were placed at the base of the subtree (Fig. 1), generating discrepancies in the gene tree topology and the phylogeny of early land plant lineages.

### Phylogenomic Synteny Network Analyses of *ARF* genes

The broad-scale phylogenetic analyses suggested some subtree topologies that are consistent with the occurrence of ancient gene duplications but genomic synteny analyses are required to provide more substantive evidence(Tang *et al.*, 2008). The recently established synteny network approach, taking advantages of accumulated plant genomes, was able to provide such substantive evidence for ancient evolutionary processes of a specific gene family(Zhao *et al.*, 2017, Zhao and Schranz, 2017). Applying this approach, we conducted a phylogenomic syntenic network analyses for *ARF* genes using a collection of available plant genomes (Supplementary Fig. S1). Syntenic *ARF* genes (syntelogs) were observed in some non-flowering plants (e.g. a lycophyte and a moss), but all represented in-paralogues which were considered to have derived from lineage-specific duplications and e *ARF* genes identified in angiosperm genomes were the primary target of the analysis and used as anchors to construct the genomic synteny network.

Among the 1,227 annotated angiosperm *ARF* genes containing valid B3 and Auxin_resp domains (Supplementary Table S2), 1,096 (89.3%) were detected to be located within genomic synteny regions that demonstrated genomic collinear relationships with at least another one *ARF* gene, and a total of 18,511 syntenic connections among *ARF* genes were detected (Figs. 3A and 3B). Consistent with the family phylogeny described previously, most of the genome syntenic connections were observed within each of the six groups. *ARF* genes from distinctive ARF groups were syntenically connected (Fig. 3A), for example, *ARF* genes from group VI were connected to *ARF* genes from group III/IV/V and group III *ARF* genes with group I. The ARF synteny network analyses uncovered a total of 82 intergroup connections (Fig. 3A and Supplementary Table S3).

**Figure 3.**
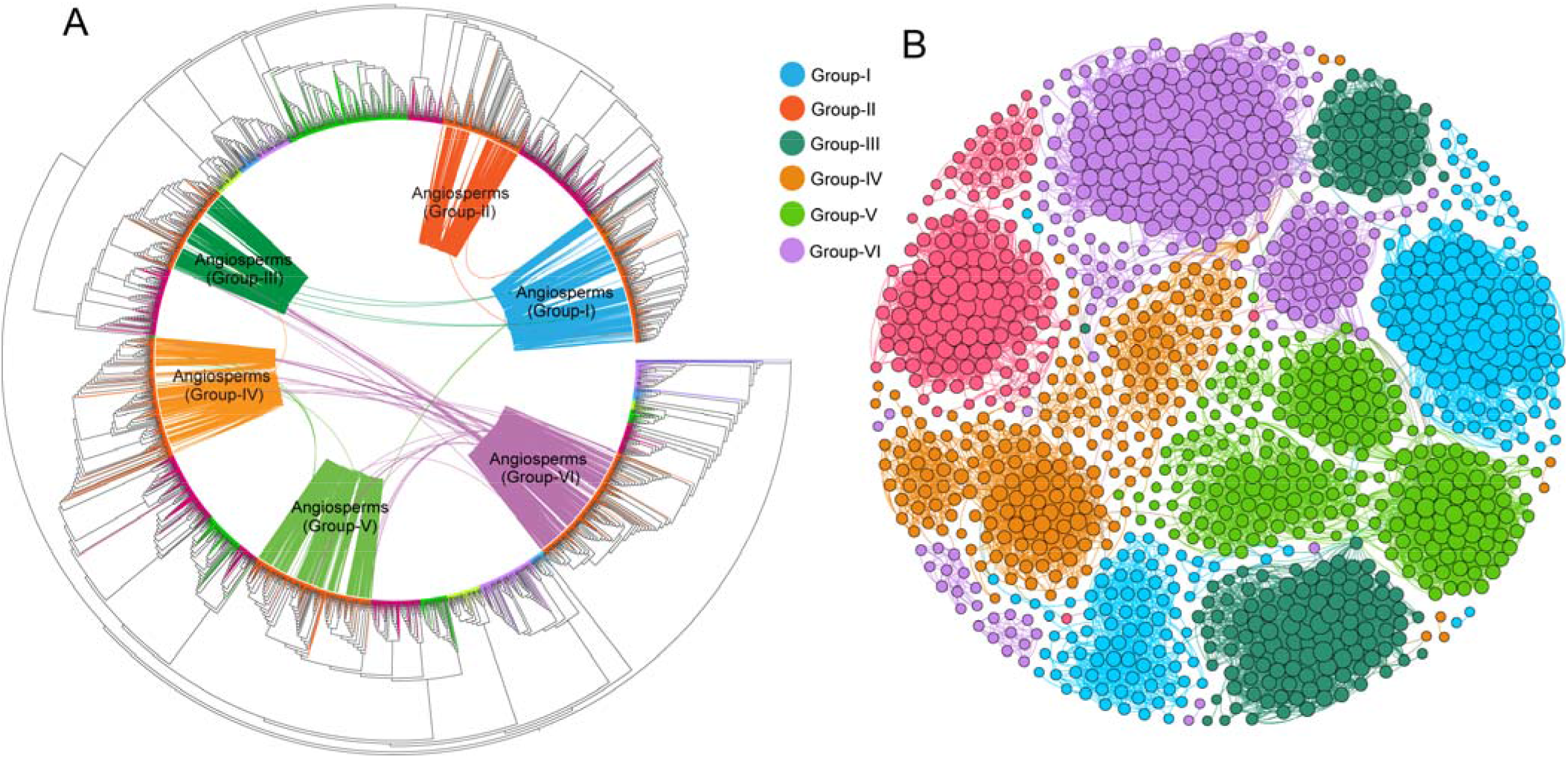
Genomic synteny analyses of ARF genes among angiosperms genomes. **(A)** Maximum-likelihood gene tree for the ARF gene family with genomic syntenic relationships between the genes. Each connecting line located inside the inverted circular gene tree (implemented in iTOL) indicates a syntenic relationship between two *ARF* genes (syntelogs). The connecting lines are colored in congruence with the six angiosperm ARF groups. **(B)** Synteny network of the ARF gene family using detected syntenic relations extracted from the genome synteny network database, using nodes representing *ARF* loci and edges (connecting lines) representing syntenic relationships. Colors of the nodes represented the six groups of *ARF* genes in angiosperms and size of each node indicates its connectivity (bigger nodes have more connections). The synteny network were clustered and visualized using the ‘Fruchterman Reingold’ layout implemented in Gephi.

In the *ARF* gene synteny network, we detected 96 *ARF* genes that did not pass our ARF identification procedure but were demonstrated to be homologous and syntenic to the annotated *ARF* genes. These syntelogs were further inspected and most contained truncated B3 and/or Auxin_resp domains or lacking either or both of these signature domains. These truncated or pseudogenes that were retained in the syntenic genomic blocks were not incorporated in the phylogenetic analyses, however, we were able to assign and label them into one of the six angiosperm *ARF* gene groups by aligning them to classified angiosperm genes. In this way, both intact (total 1,096) and truncated (total 96) *ARF* genes involved in the synteny network were classified. The classification for these truncated genes were considered reliable because of the distant phylogenetic relationships among the six groups (Fig. 1). This may suggest that using genomic syntenic relationships could be a robust approach for detecting pseudogenes retained in the syntenic genomic blocks and which exhibit significant local sequence identity with intact functional paralogues.

*ARF* genes from each group were conspicuously found in separate and distinct syntenic communities in the initial synteny network visualization (Fig. 3B). The ARF synteny network was further dissected to find subnetwork communities by the use of clique percolation clustering at k = 3 implemented in CFinder v2.0.6 (Adamcsek *et al.*, 2006). A total of 25 communities (numbered 0 through 24) (nodes clustered within a subnetwork usually possess more connections in its community than with nodes in other communities) were obtained (Fig. 4). Among the 1,192 ARF syntelogs that were extracted from the synteny network database, 1,128 (94.6%) were identified in the 25 network communities, other syntelogs that had a single syntenic connection or were not involved in a clique (at k=3) were excluded. For example, among the 22 *ARF* genes in *Arabidopsis thaliana,* 17 members were clustered in 13 synteny network communities (Fig. 4A). The chromosome-level genome assemblies represented the best material for genome synteny analyses, but some plant genome assemblies currently available are still highly fragmented. For example, in the *Malus domestica* (apple) genome, only one *ARF* gene was clustered in the synteny network because the genome assembly version we obtained from Phytozome database and that was used in our synteny network construction was fragmented (approximately 881.3 Mb arranged in 122,107 scaffolds) (Fig. 4A). However, the network approach using multiple plant genomes appeared to be error-tolerant and the results were unaffected by the inclusion of a few fragmented genomes(Zhao *et al.*, 2017).

**Figure 4.**
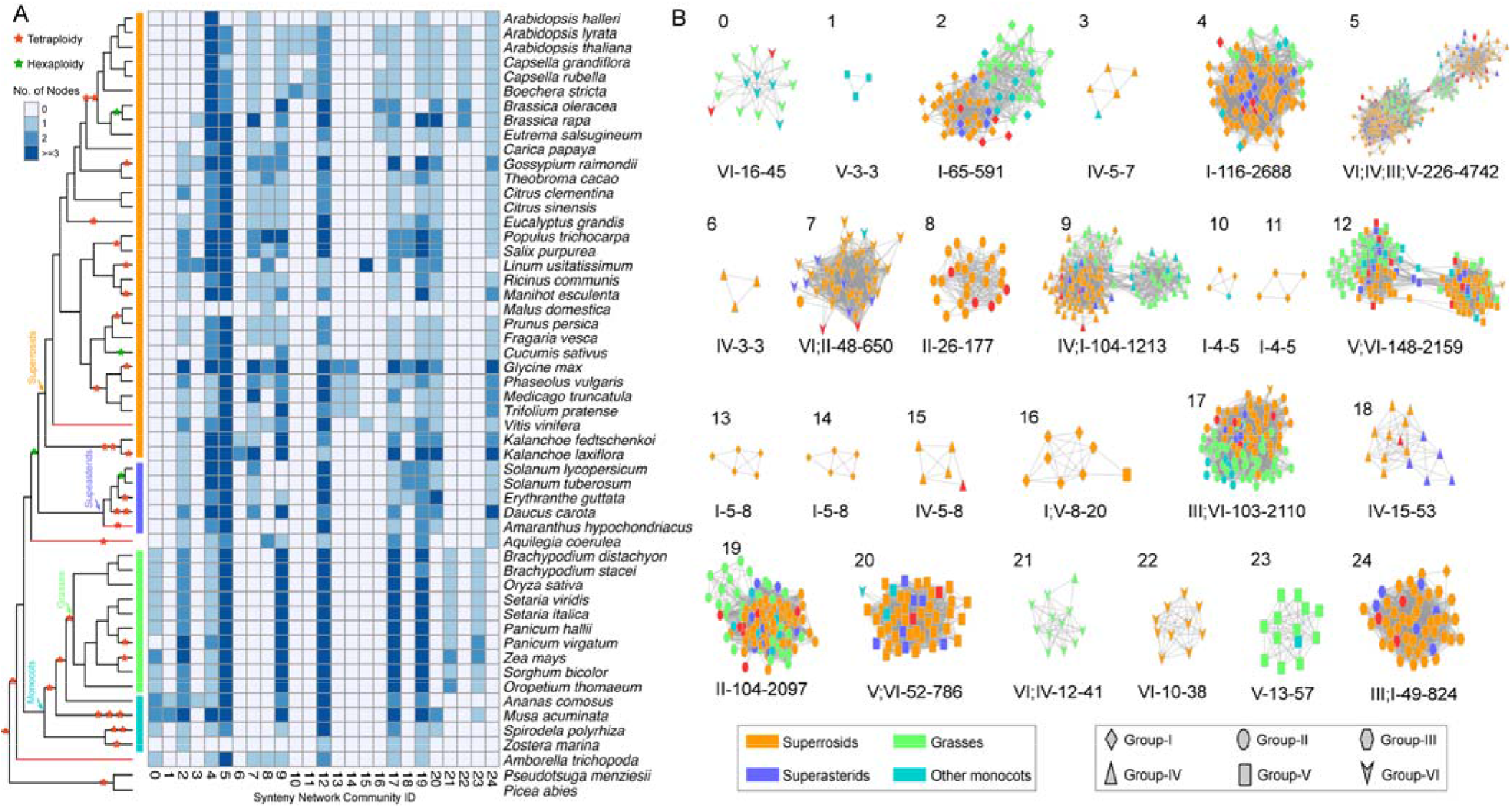
Detailed network representations for ARF synteny network communities among angiosperm genomes. **(A)** Species composition for each of the 25 network communities. Blue-colored cells depict the presence of ARF syntelogs in the different species. The 25 network communities were identified using CFinder at k=3. **(B)** Detailed visualization for each of the ARF synteny network communities. Nodes in different colors represented different plant lineages, and the node shapes represented different ARF subfamilies (Groups I through VI). Selected basal lineages including *Vitis vinifera* (sister to other rosids), *Amarsanthus hypochondriacus* (sister to asterids), and *Amborella trichopoda* (the basal angiosperm) were depicted as red nodes in the communities. Each network community was presented following the ‘ARF clades’-‘number of nodes’-‘number of connections’ nomenclature system and some communities contain genes from multiple ARF clades. One synteny community may contain genes from various groups.

Species compositions for each of the 25 synteny network communities (Fig. 4A) indicate that network communities 4 and 5 are angiosperm-wide, containing *ARF* genes from monocots, eudicots and *Amborella,* Community 23, on the other hand, only contains *ARF* genes from monocots and community 24 is solely confined to *ARF* genes from eudicots. Other communities are lineage specific such as community 21 which only contains *ARF* genes from grasses, communities 13 and 14 that are specific to legumes, and communities 16 and 22 that are specific to the Brassica.

Subnetwork communities were separately visualized, using node colors to depict different plant lineages and node shapes, to delineate *ARF* genes from the different classification groups (Fig. 4B). Community 0 (labeled as ‘VI-16-45’) consisted of *ARF* members from group-VI, with a total of 16 nodes and 45 connections within the community. Some syntenic communities contained *ARF* genes from multiple groups. Community 5 was recognized as the largest community with 226 nodes and 4,742 connections, and nodes in this community were primarily *ARF* genes from group-VI and group-IV, with a minority of members from group-III (3 nodes) and group-V (1 nodes). The mixed group communities suggest the existence of ancient tandem duplications(Zhao *et al.*, 2017), where duplicated paralogues were likely lost in the ancestral genome such that ancient tandem paralogues are not seen in most current plant genomes, but synteny network analyses reflect them as multigroup communities. Consistent with this hypothesis, tandem *ARF* genes from distant groups were not present in the genome of a single species used in the analysis (Supplementary Table S1). To illustrate this, the *ARF* gene (scaffold00029187) from *Amborella* was classified as a group-IV member, but it had a syntenic connection with group-VI *ARF* genes from *Oryza sativa* (LOC_Os10g33940), *Oropetium thomaeum* (Oropetium_20150105_02810A) and *Phaseolus vulgaris* (Phvul_003G075800). This could be explained by the occurrence of an ancient tandem gene duplication that was generated prior to the separation of groups VI and IV. Following the speciation of basal angiosperms and eudicots plus monocots, the group-VI member was lost in *Amborella*, and the group-IV member was lost in the ancestor of monocots and eudicots resulting in the syntenic relationship seen between group-VI and group-IV *ARF* genes. The inter-group genomic syntenic connections not only provided evidence for ancient gene duplications followed by lineage-specific gene losses, but also suggested that modern *ARF* genes evolved from a common ancestor present in the streptophytes.

### Evolutionary characteristics for each of the six groups of ARFs in angiosperms

The global phylogenetic and synteny network analyses generated a robust six-group classification system for *ARF* genes and indicated pervasive intra-group syntenic phylogenetic relationships. To elaborate the evolutionary processes within each of the six groups, individual phylogenetic trees for angiosperm genes in each of the six groups were estimated separately and syntenic connections within each network community were mapped onto the six gene trees (Gamboa-Tuz *et al.*, 2018). Along with the *ARF* geness identified from Phytozome plant genomes, the *ARF* genes from basal angiosperms (also ANA grade) and magnoliids were incorporated in the phylogenetic analyses, however these *ARF* gene sequences were derived from transcriptomes and thus did not provide syntenic information. The number of angiosperm (including eudicots, monocots, magnoliids and ANA grade) *ARF* genes in each of the six groups ranged from 190 (group II) to 318 (group I). Below we describe the primary evolutionary characteristics for the six ARF groups separately.

#### Group-I

This group represented the largest clade (containing 318 angiosperm *ARF* gene members) of the six groups (Fig. 5A). An evident angiosperm-wide duplication (delineated as groups IA and IB) was identified from the tree topology with the three relevant bootstrap values supporting the duplication node and the two child clades greater than 95%. Both IA and IB clades include genes from monocots, eudicots, magnoliids and basal angiosperm lineages. The single *ARF* gene member from *Amborella* was placed as sister to the IA plus IB duplication clade, suggesting that the *ARF* gene duplication likely occurred after the separation of *Amborella* from other angiosperms.

**Figure 5.**
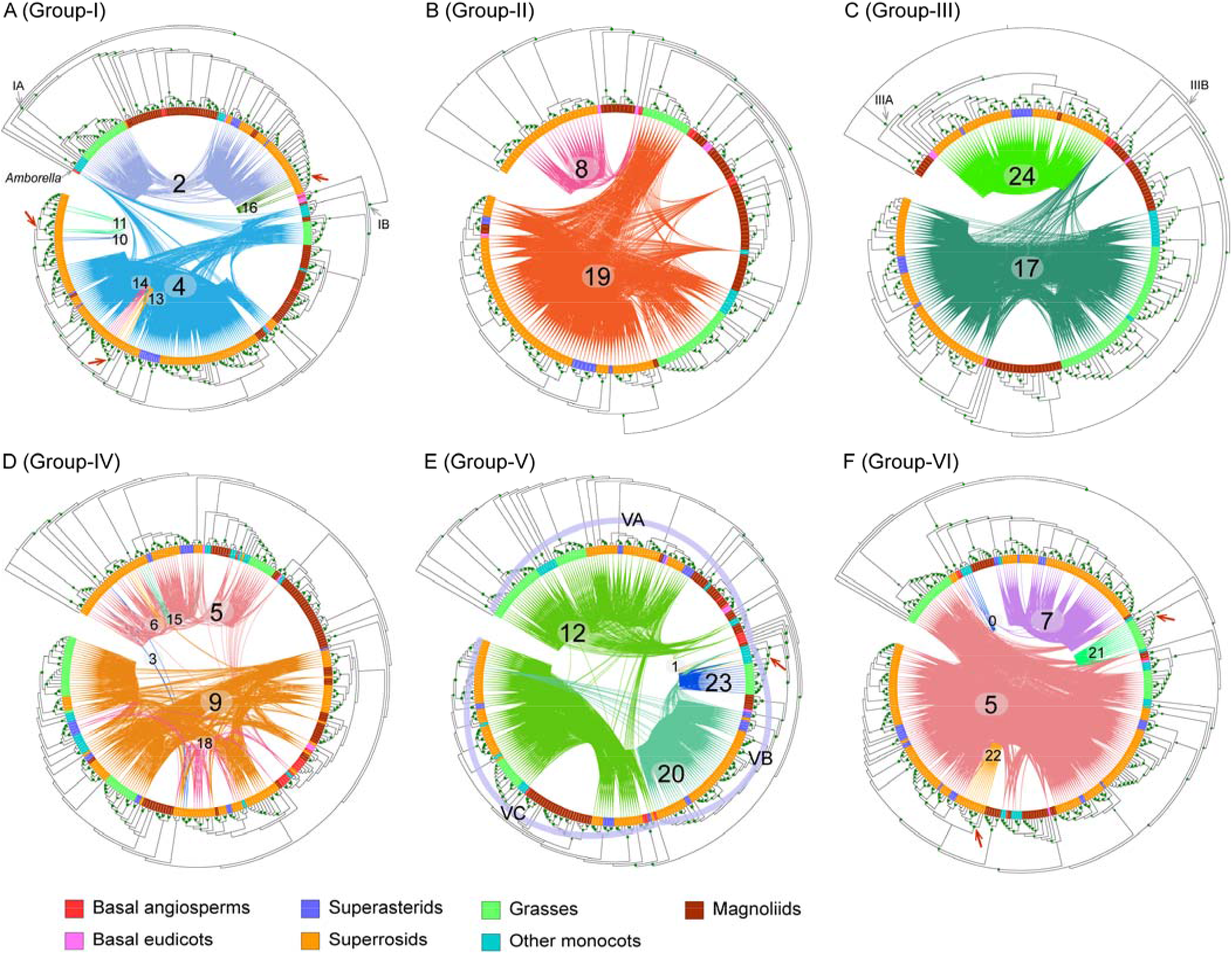
Phylogenetic and synteny network analyses for each of the six groups of ARFs in angiosperms. Maximum-likelihood trees for each of the six ARF groups were constructed, genes from different species groups were colored using different colors and genes detected in syntenic genomic blocks (syntelogs) were connected using curved lines. The syntenic connections belonging to different synteny network communities were plotted using different colors. Synteny network communities were numbered according to that depicted in figure 2B. Inferred ancestral transposition activities were indicated by red arrows.

Network communities associated with group-I primarily included angiosperm-wide communities 4 (116 nodes) and 2 (65 nodes) (Fig. 4B), which align to groups IA and IB (Fig. 5A), respectively. Group IA was consistent with the designation ARF9 and group IB and the designation ARF1 in *A. thaliana* as reported previousely(Finet *et al.*, 2013). The number of *ARF* genes included in group IA was conspicuously greater than in group IB, particularly for the *ARF* genes from superrosids. The core-eudicot duplication (also known as gamma event)(Jiao *et al.*, 2012) may have contributed to the family expansion, but some ARF genes from magnoliids were also included in the duplication clade and the bootstrap supporting value for the duplication node was also lower than 70%. Moreover, some lineage-specific network communities for *ARF* genes in group-I were observed, where communities 10, 11, 13, 14 and 16 are small communities containing the *ARF* genes from superrosids (Fig. 4B) and these syntenic communities rendered as monophyletic clades in the phylogenetic analyses (Fig. 5A). The species composition analysis (Fig. 4A) for these lineage-specific communities indicated ancestral transposition activities in the Brassicaceae (communities 10, 11 and 16) and legumes (communities 13 and 14).

#### Group-II

Group-II was the smallest group (containing 190 angiosperm *ARF* genes) and synteny network analyses revealed two primary communities, 19 and 8, as depicted in Fig. 5B. Community 8 contains 26 nodes with *ARF* genes from only eudicots and *Amborella*, clustered with a group of magnoliid genes, that formed a paraphyletic clade at the basal position. While the nodes in community 19 were angiosperm-wide, and *ARF* genes from grasses were separated into two clades, one clade following the *ARF* genes in community 8 and the other clade clustered with the other monocots. However, the *ARF* genes clustered in each of the two grass clades did not share syntenic connections (Fig. 5B), and the two basal species (*Aquilegia* and *Amborella)* were included in both communities (Fig. 4B), suggesting the genome context (e.g. regulatory elements and adjacent genes) were altered for the *ARF* genes in the two communities.

The nodes clustered in community 19 may correspond to an ancient tandem duplication in the ancestor of angiosperms as a clade of *ARF* genes from the grasses were evidently separated from other nodes in community 19, indicative of more intra than inter connections (Fig. 4B).

#### Group-III

Group-III contains 216 *ARF* genes incorporated primarily in network communities 17 and 24 in the synteny network analyses (Fig. 4B and Fig. 5C). The phylogenetic profile for group-III genes identified them as forming two well-separated monophyletic clades (delineated as IIIA and IIIB in this study). The group-IIIA (community 24) contains *ARF* genes from only eudicots and magnoliids, while community 17 is angiosperm-wide and recognized as group-IIIB. The species composition analysis of group-IIIA encompassed a core-eudicot duplication (gamma event), although a magnoliids *ARF* gene was also in this group, that was shared by superrosids and superasterids. *ARF* genes from basal eudicots are recognized as sister to this duplication clade. Similarly, a duplication clade shared by *ARF* genes from grasses (and one gene from pineapple) was conspicuous and likely contributed to the generation of more *ARF* gene members in group-IIIB in the grasses.

#### Group-IV

Group-IV contains 282 angiosperm *ARF* genes that were contained in six major network communities 5, 9, 18, 15, 3 and 6; community 5 was the largest community containing genes from multiple groups (Fig. 4B). Network communities 5 and 9 are angiosperm-wide and 18 contains *ARF* genes from only eudicots. The remaining three communities (3, 6 and 15) were smaller and none formed a high-confidence monophyletic clade which in turn does not support the possibility of ancestral lineage specific transpositions. By comparing the genomic synteny connections with the phylogenetic profile, two evident clusters of *ARF* genes within this group were recognized (Fig. 5D). Community 5 (also communities 6 and 15) was clustered into one group and communities 9 and 18 were clustered into another. Both groups were recognized as angiosperm-wide groups suggesting an angiosperm-wide duplication within group-IV, athough the duplication topology cannot be easily deduced from the gene tree. In the community 9 network, a subnetwork of monocot *ARF* genes contained more intra-connections than other communities and were separated from other nodes (Fig. 4B), suggesting the possibility of extra rounds of gene duplications and losses in the evolutionary past of *ARF* genes in group IV in angiosperms.

#### Group-V

Group-V contains a total of 287 angiosperm *ARF* genes that were clustered in four synteny communities, 12, 20, 23 and 1 (Fig. 5E), among which communities 12 and 20 were angiosperm-wide, and communities 23 and 1 contain small numbers of monocots *ARF* genes. By integrating the synteny network and phylogenetic profile analyses, three subgroups could be identified (delineated as VA, VB and VC), and consistent with the community network analyses, the nodes in community 12 were phylogenetically separated into two subgroups (groups-VA and -VC). Nodes in communities 23 and 1 were recognized in one monophyletic clade (group-VB). An ancestral transposition in the ancestor of commelinids (including grasses, pineapple and banana genes, community 23 and community 1) (Figs. 4A and 4B), and an *ARF* gene from *Spirodela polyrhiza* (duckweed, sister to commelinids) was syntenically clustered in community 20.

#### Group-VI

The Group-VI included 295 *ARF* genes integrated into 5 network communities 5, 7, 0, 21 and 22 (Fig. 5F). Community 5 was conspicuously angiosperm-wide, community 7 encompassed primarily *ARF* genes from eudicot and *Amborella,* community 0 contained *ARF* genes from monocots and *Amborella,* and communities 21 and 22 were solely comprised of *ARF* genes from grasses and crucifers, respectively (Fig. 4A). Mapping the syntenic connections on the phylogenetic tree, conspicuous monophyletic clades in grasses (community 21) and crucifers (community 22) were generated and provided phylogenomic evidence for ancestral transposition activities in these two lineages. The *ARF* genes clustered in community 5 were phylogenetically separated into two distinct clades with some *ARF* genes from grasses were placed in a basal position in the group-VI phylogeny. The nodes in community 7 were well-clustered in the family phylogeny.

In the phylogenomic synteny network analyses we employed the maximum-likelihood gene tree generated by IQ-TREE in which more evolutionary models were implemented. We attempted to reconstruct the *ARF* gene family phylogeny using RAxML and the PROTGAMMAAUTO model (Supplementary Fig. S3), which generated alternative tree topologies, nevertheless, the syntenic community patterns remained steady and the major duplication clades and transposition activities could be consistently captured. Tree uncertainties may make some of the evolutionary processes of *ARF* gene family elusive, but the synteny network approach appears robust and uncovered evolutionary details and provided more clues for future experimental studies. For example, *ARF* genes were recurrently duplicated and transposed in specific lineages which suggests that the functions of these transposed genes might reveal novel regulatory elements that were captured in their altered genomic context. The transpositions that we indicate to have occurred in crucifers, legumes, commelinids and grasses were tightly associated with ancestral polyploidy events(Van de Peer *et al.*, 2017), which generated more possibilities in the gene regulatory network. The ancestral gene duplication together with transpositions could have greatly contributed to the expansion of the auxin regulatory network which would have had important implications in the understanding of the evolutionary processes of current land plants.

## CONCLUSION

In this study, we generated a broad-scale family phylogeny for *ARF* genes from augmented genome and transcriptomic data, that updated our current understanding of the evolutionary history of this transcription factor in streptophytes. Based on the family phylogeny, we proposed a six-group classification regime for angiosperm *ARF* genes. Group IV contains the ARF activators and these genes are well-conserved in all land plant clades. The Group IV subfamily phylogeny also supported the ‘hornwort-sister’ hypothesis. Genomic synteny network analyses revealed highly-conserved genomic syntenies among angiosperm *ARF* gene loci and within each of the six *ARF* gene groups. CFinder clique analyses of the *ARF* gene synteny network identified 25 subnetwork communities, which were further projected onto the six subfamily phylogenies. The analyses suggest that ancient duplications and transpositions have greatly contributed to the diversification of *ARF* genes in angiosperms. Ancestral lineage-specific transpositions involving *ARF* genes were unveiled in crucifers, legumes, commelinids and grasses in groups I, V and VI. Future studies focusing on non-angiosperm specific lineages should benefit from the evolutionary framework used in this study, especially when more genomes in these plant lineages become available(Cheng *et al.*, 2018).

## MATERIALS AND METHODS

### Collection of Auxin Response Factors

To generate a broad-scale family phylogeny homologues of plant *ARF* transcription factor genes were obtained from Phytozome v12.1.6 (https://phytozome.jgi.doe.gov/pz/portal.html) and the OneKP (https://db.cngb.org/onekp/)(Matasci *et al.*, 2014) databases using blastp searches filtered with an e-value threshold of 1e-5. *ARF* gene sequences from fern genomes were collected from FernBase (https://www.fernbase.org)(Li *et al.*, 2018). The protein domain composition of each of the putative ARF protein sequences were determined by querying the NCBI Conserved Domain Database(Marchler-Bauer *et al.*, 2017) and only sequences that contained both definitive functional domains: B3 DNA-binding domain (Pfam accession: PF02362) and Auxin_resp (Pfam accession: PF06507), were included in subsequent analyses (Supplementary Table S2).

### Family Phylogeny Construction

To generate reliable sequence alignments for the collected *ARF* gene-family members, boundaries of the B3 and Auxin_resp domains were identified by aligning each of the protein sequences onto the two HMM profiles using hmmalign v3.2.1(Eddy, 2008, Eddy, 2011). Alignments of the two domains were separately refined using muscle v3.8.1551 and concatenated to generate a broad-scale sequence alignment for *ARF* genes. Columns in the alignment with more than 20% gaps were removed using Phyutility v2.2.6(Smith and Dunn, 2008).

IQ-TREE v1.6.8(Nguyen *et al.*, 2015) software was employed to reconstruct the maximum likelihood (ML) gene tree. For the obtained broad-scale amino acid alignment, the JTT+R9 model was the best-fit evolutionary model selected by ModelFinder(Kalyaanamoorthy *et al.*, 2017) under Bayesian Information Criterion. The SH-aLRT test and ultrafast bootstrap (Hoang *et al.*, 2018) analyses with 1000 replicates were conducted in IQ-TREE to obtain the supporting values for each internal node of the tree. The obtained maximum-likelihood gene trees were visualized and edited using FigureTree v1.4.4 (http://tree.bio.ed.ac.uk/software/figtree/) and iTOL v4.3 (https://itol.embl.de)(Letunic and Bork, 2016). Maximum-likelihood trees for each of the six angiosperm *ARF* clades using IQ-TREE (including model-selection procedure) were also reconstructed to infer potential duplication nodes by analyzing the detailed clade-specific phylogenies.

The phylogenetic analyses for each of the six ARF groups were performed using both IQ-TREE v1.6.8 (Nguyen *et al.*, 2015) and RAxML v8.2.12 (Stamatakis, 2014). The model selection procedure was performed within IQ-TREE based on the Bayesian information criterion (BIC) and for RAxML analyzes we used the ‘-m PROTGAMMAAUTO’ model with 500 bootstrap replicates. All trees were inspected, but the IQ-TREE algorithm produced better bootstrap support overall for branches (Fig. 5 and Supplementary Fig. S3). Each of the six ARF groups contained multiple synteny network communities and syntenic connections in different communities were plotted using different colors as implemented in the iTOL v4.3(Letunic and Bork, 2016).

### Genomic synteny network construction

To unveil the genomic syntenic relationships among plants, protein sequences for each of the 52 angiosperm genomes were compared with each other and themselves using Diamond v0.9.22.123 software(Buchfink *et al.*, 2015) with an e-value cutoff at 1e-5. In this way, blastp tables for a total of 52×51/2+52=2,704 whole proteome comparisons were generated. Only the top five non-self blastp hits were retained as input for the MCScanX(Wang *et al.*, 2012a) analyses. The *ARF* gene associated syntenic genomic block were extracted (Supplementary Table S3) and visualized in Cytoscape v3.7.0(Shannon *et al.*, 2003) and Gephi v0.9.2(Bastian *et al.*, 2009). Some ARF syntelogs were truncated or demonstrate absence of signature domains and were not included in our phylogenetic analyses. These truncated ARF genes were classified and labelled (clade I through VI) by comparing with those classified as *ARF* genes. The phylogeny of angiosperm species and the associated paleopolyploidy events were redrawn based on a tree reported earlier by Van de Peer *et* al.(Van de Peer *et al.*, 2017) and the APG IV system(Byng *et al.*, 2016) with minor modifications: the hexapolyploidy event in cucurbitaceae(Wang *et al.*, 2018), the fern genome duplications(Li *et al.*, 2018), the ancestral duplication events mosses(Devos *et al.*, 2016, Lang *et al.*, 2018) and in Caryophyllales(Yang *et al.*, 2018), were included in the tree.

The ARF syntenic networks were analyzed using CFinder v2.0.6(Adamcsek *et al.*, 2006) utilizing the unweighted CPM algorithm and no time limit. All possible k-clique (from 3 to 21) communities were identified for the complete *ARF* gene syntenic network. We used k=3 as the clique community threshold and in this scenario one *ARF* gene (node) involved in a subnetwork community should have at least two connections (edge) with other nodes in the community. Increasing k values made the communities smaller and more disintegrated but also more connected. For illustration purposes, we used different nodal shapes to represent the members from the six ARF groups and different colors to depict specific plant lineages using the Cytoscape v3.7.0 software(Shannon *et al.*, 2003). For each of the 25 communities, the species composition of the syntelogs were counted and a heatmap was generated using the pheatmap v1.0.10 (https://github.com/raivokolde/pheatmap) package implemented in the R statistical environment.

## ACKNOWLEDGEMENTS

This work was supported by Hong Kong Research Grant Council (GRF CUHK12100318, AoE/M-05/12, AoE/M-403/16), the Fundamental Research Funds for the Central Non-profit Research Institution of CAF (No. CAFYBB2018QB002 to L.W.) and the National Science Foundation (Collaborative Research: Dimensions: No. 1638972 to M.J.O.).

## AUTHOR CONTRIBUTIONS

BG conceived the study, performed the bioinformatic analyses and wrote the manuscript. XL and MC contributed to the data collection and discussion. JZ critically revised and improved the manuscript. All authors read and approve the final manuscript.

## SUPPORTING INFORMATION

Additional Supporting Information may be found in the online version of this article.

**Figure S1**. Plant lineages screened for ARF homologues.

**Figure S2**. Number of Auxin Response Factor genes identified from each of the plant genomes.

**Figure S3**. Phylogenic and synteny network analyses for each of the six groups of ARFs in angiosperms.

**Table S1**. Annotation and Classification of ARF genes in *Arabidopsis thaliana.*

