## Supplementary figures & table S1 for "Evolution of Auxin Response Factors in plants characterized by phylogenomic synteny network analyses"

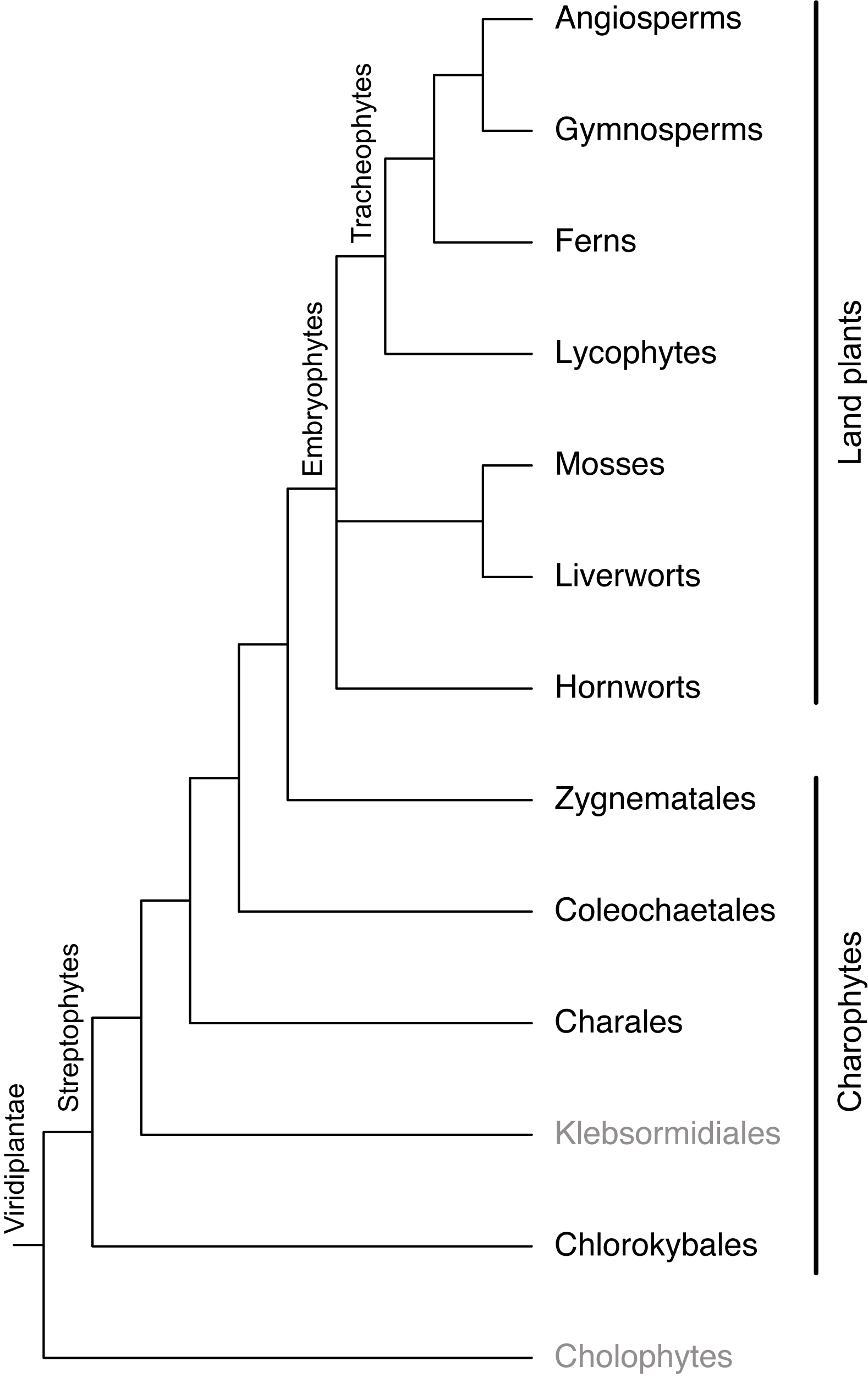

**Supplementary Figure S1.** Plant lineages screened for ARF homologues. Auxin response factors were not detected in lineages indicated in grey. Relationships for plant lineages were redrawn from (Wilhelmsson et al., 2017) with minor modifications.

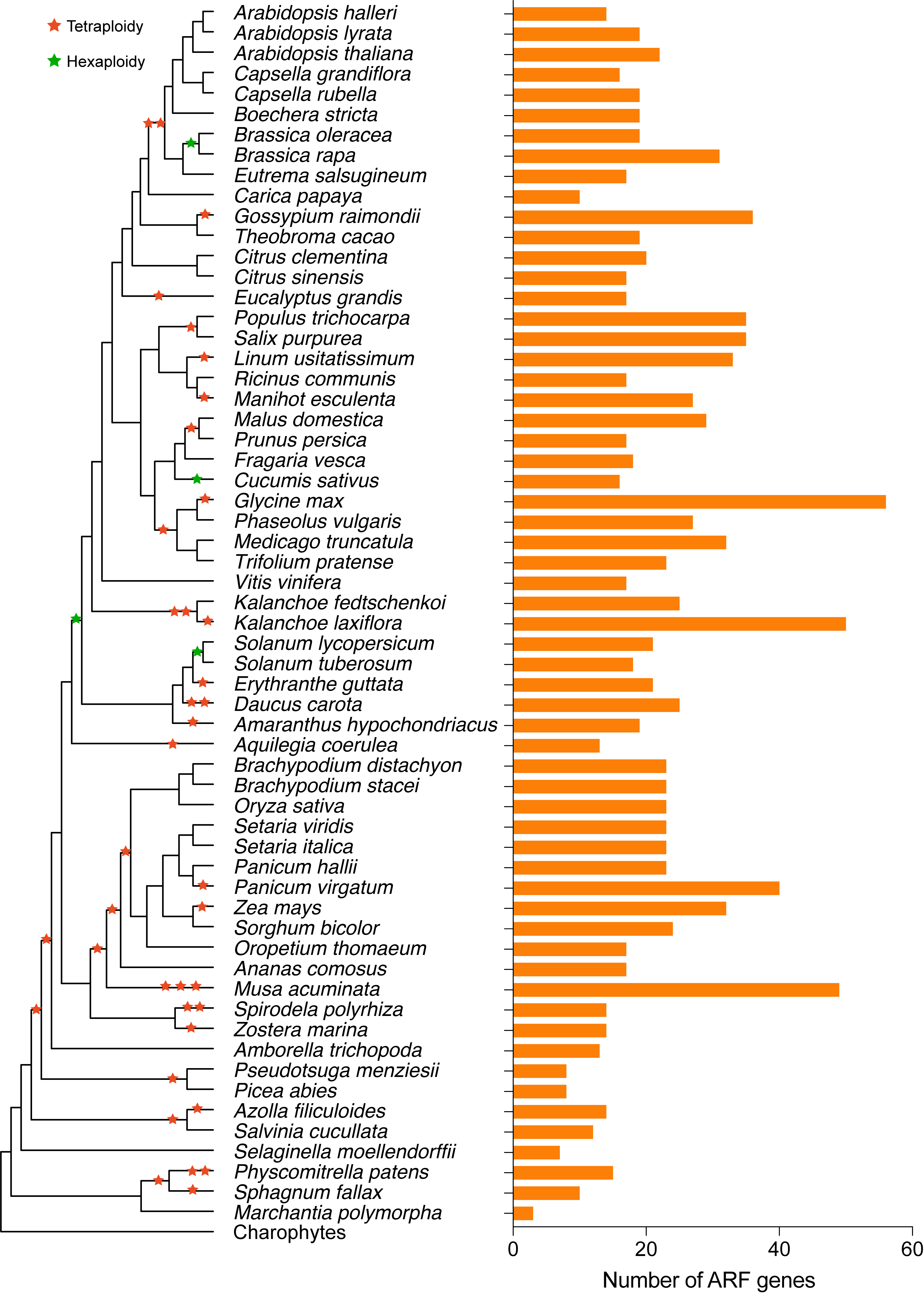

**Supplementary Figure S2.** Number of Auxin Response Factor genes identified from each of the plant genomes. The species tree and related paleo-polyploidy events was plotted according to (Van de Peer et al., 2017) with minor modifications.

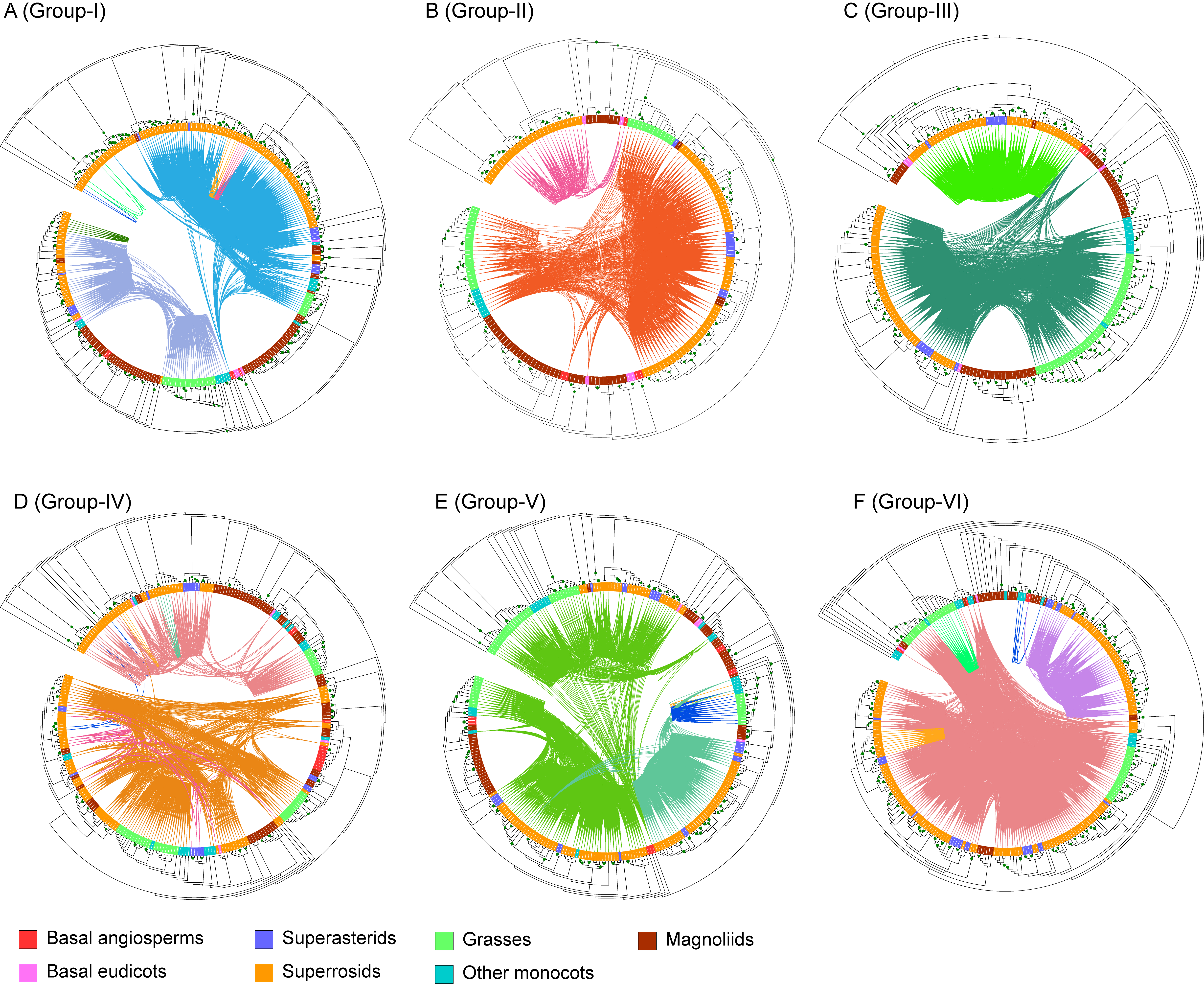

**Supplementary Figure S3.** Phylogenic and synteny network analyses for each of the six groups of ARFs in angiosperms. Maximum-likelihood trees (constructed using RAxML) for each of the six ARF groups were constructed, genes from different species groups were colored using different colors and genes detected in syntenic genomic blocks (syntelogs) were connected using curved lines. The syntenic connections belonging to different synteny network communities were plotted using different colors.

**Supplemental Table S1** – Annotation and Classification of *ARF* genes in *Arabidopsis thaliana*

| Gene ID | TAIR description | Annotation | Finet *et al.* 2013 | This Study |
| --- | --- | --- | --- | --- |
| AT1G34170 | auxin response factor 13 | ARF13 | ARF 9 | Group_I |
| AT1G34310 | auxin response factor 12 | ARF12 | ARF 9 | Group_I |
| AT1G34390 | auxin response factor 22 | ARF22 | ARF 9 | Group_I |
| AT1G34410 | auxin response factor 21 | ARF21 | ARF 9 | Group_I |
| AT1G35240 | auxin response factor 20 | ARF20 | ARF 9 | Group_I |
| AT1G35520 | auxin response factor 15 | ARF15 | ARF 9 | Group_I |
| AT1G35540 | auxin response factor 14 | ARF14 | ARF 9 | Group_I |
| AT1G59750 | auxin response factor 1 | ARF1 | ARF 1 | Group_I |
| AT2G46530 | auxin response factor 11 | ARF11 | ARF 9 | Group_I |
| AT3G61830 | auxin response factor 18 | ARF18 | ARF 9 | Group_I |
| AT4G23980 | auxin response factor 9 | ARF9 | ARF 9 | Group_I |
| AT5G62000 | auxin response factor 2 | ARF2 | ARF 2 | Group_II |
| AT2G33860 | auxin response factor 3 | ARF3 | ARF 3/4 | Group_III |
| AT5G60450 | auxin response factor 4 | ARF4 | ARF 3/4 | Group_III |
| AT1G30330 | auxin response factor 6 | ARF6 | ARF 6/8 | Group_IV |
| AT5G37020 | auxin response factor 8 | ARF8 | ARF 6/8 | Group_IV |
| AT1G19220 | auxin response factor 19 | ARF19 | ARF 5/7 | Group_V |
| AT1G19850 | auxin response factor 5 | ARF5 | ARF 5/7 | Group_V |
| AT5G20730 | auxin response factor 7 | ARF7 | ARF 5/7 | Group_V |
| AT1G77850 | auxin response factor 17 | ARF17 | ARF 10/16/17 | Group_VI |
| AT2G28350 | auxin response factor 10 | ARF10 | ARF 10/16/17 | Group_VI |
| AT4G30080 | auxin response factor 16 | ARF16 | ARF 10/16/17 | Group_VI |
| AT1G43950* | auxin response factor 23 | ARF23 | ARF 9 | Group I |

*The ARF23 in *Arabisopsis thaliana* is a truncated gene.
